# Intrathecal administration of bone marrow stromal cells and TGF-β1 alleviate chemotherapy-induced neuropathic pain in male mice

**DOI:** 10.1101/2022.10.19.512871

**Authors:** Yul Huh, Xin Luo, Di Liu, Changyu Jiang, Ru-Rong Ji

## Abstract

Chemotherapy-induced peripheral neuropathy (CIPN) is the de facto clinical side effect that limits the administration of anti-cancer treatments. Recently, we reported that intrathecally injected bone marrow stromal cells (BMSCs) reduced nerve trauma-induced neuropathic pain in male mice via TGF-β1 signaling. In this study, we examined sex-dependent pain relief mediated by intrathecally delivered BMSCs and TGF-β1 in paclitaxel (PTX)-induced CIPN. BMSCs were prepared from primary cultures of male or female mice separately. A single intrathecal injection of BMSCs, prepared from male donors, completely prevented the development of PTX-evoked mechanical allodynia in male mice. However, female mice showed no analgesic response to either male or female BMSCs. Additionally, male mice did not demonstrate an analgesic response to BMSCs from female donors. Intrathecal injection of TGF-β1 neutralizing antibody reversed the analgesic action of BMSCs. Interestingly, spinal administration of TGF-β1 reduced mechanical allodynia in male mice but not in female mice. Ex vivo patch-clamp recordings in spinal cord slices revealed that TGF-β1 inhibited PTX-induced synaptic plasticity, i.e. increase in spontaneous excitatory synaptic currents (sEPsCs), in spinal cord neurons from male mice only. Intrathecal TGF-β1 increased the paw withdrawal threshold in von Frey testing in naïve mice of males but not females, and the antinociceptive effect of TGF-β1 in males was blocked by orchiectomy-induced androgen deficiency. Together, these findings reveal sex dimorphism in BMSC control of mechanical pain through spinal TGF-β1 signaling.

## Introduction

Sex-dimorphism in chronic pain is a well-characterized clinical phenomenon, as more women suffer from pathological pain such as headaches, migraine, and fibromyalgia, than men (Riley et al., 1998; Bartley and Fillingim, 2013; Navratilova et al., 2021). However, most of the preclinical pain research have been conducted in male animals, leading to a biased literature in pain research (Mogil, 2012, 2020). Recent progress in preclinical research has demonstrated considerable sex differences in neuroimmune interactions in pathological pain. For example, inhibition of spinal microglial signaling cascades of toll-like receptor 4 (TLR4), p38 mitogen-activated protein kinase (MAPK), and brain-derived neurotrophic factor (BDNF) reduced neuropathic pain and inflammatory pain in male but not female animals (Sorge et al., 2011; Sorge et al., 2015; Taves et al., 2016; Chen et al., 2018; Luo et al., 2018; Mapplebeck et al., 2018; Agalave et al., 2021). In female animals, T cells are thought to mediate pathological pain (Sorge et al., 2015; Rosen et al., 2017). Interestingly, regulatory T-cells were shown to inhibit microglial activities in the spinal cord of female mice (Kuhn et al., 2021). Accumulating evidence has also shown that macrophages regulate pain in a sex-dependent manner (Luo et al., 2019; Yu et al., 2020; Luo et al., 2021).

Chemotherapy-induced peripheral neuropathy (CIPN) is a neurological disorder caused by chemotherapeutic reagents such as paclitaxel (PTX) (Forsyth et al., 1997; Sisignano et al., 2014). CIPN has been shown to cause premature termination or reduced dosage of chemotherapy regimens, affecting survival rates of cancer patients (Seretny et al., 2014). CIPN-induced nociceptor hyperexcitability was demonstrated in both rodent and human dorsal root ganglion (DRG) sensory neurons (Li et al., 2015; Chang et al., 2018; Li et al., 2018). While chemotherapy produces limited microgliosis in the spinal cord (Zheng et al., 2011; Zhang et al., 2012), it results in remarkable recruitment of immune cells, including macrophages and T cells to the DRG (Liu et al., 2014; Zhang et al., 2016; Luo et al., 2019). Depletion of macrophages alleviated neuropathic pain in chemotherapy-treated animals (Zhang et al., 2016). However, the role of mesenchymal stromal/stem cells (MSC) or bone marrow stem/stromal cells (BMSCs) in CIPN is not fully investigated, although nasal administration of MSC was shown to reverse CIPN in mice (Boukelmoune et al., 2021).

Recently, we demonstrated that intrathecal injection of BMSCs reduced nerve trauma-induced neuropathic pain for more than two months in two mouse models of neuropathic pain, induced by chronic constriction injury and spared nerve injury (Chen et al., 2015). We also found that transforming growth factor beta-1 (TGF-β1) was the factor secreted by BMSCs and responsible for neuropathic pain relief in male mice (Chen et al., 2015). In this study, we tested the sex dimorphism of BMSC-produced pain relief in the paclitaxel model of CIPN. We also investigated the involvement of TGF-β1 signaling in paclitaxel-induced neuropathic pain and spinal cord synaptic transmission in male and female mice. Our results revealed a male-specific contribution of BMSCs and TGF-β1 signaling to CIPN and mechanical pain.

## Materials and methods

### Animals and chemotherapy-induced peripheral neuropathic pain model

Adult male and female CD1 mice (8-12 weeks old) were purchased from Charles River Laboratories. Animals were randomly assigned to each group. All animals were maintained by the Division of Laboratory Animal Resources (DLAR) at the Duke University Animal Facility. All animal experiments were approved by the Institutional Animal Care and Use Committees (IACUC) of Duke University. CIPN was induced in mice by 4 intraperitoneal injections of paclitaxel (Sigma-Aldrich) at dose of 2 mg/kg on days 1, 3, 5, and 7 (Luo et al., 2018).

### Drugs and administration

TGF-β1, and TGF-β1 neutralizing antibody, and TGF-β1 ELISA kit were purchased from R&D Systems. Paclitaxel was purchased from Sigma-Aldrich.

### Intrathecal injection

Drug or cells were delivered to cerebrospinal fluid by lumbar puncture, which can target both DRG and spinal cord (Hylden and Wilcox, 1980; Kawasaki et al., 2008) via intrathecal (I.T.) route. Mice were briefly anesthetized with isoflurane (2%) and a lumbar puncture was performed between the L5-L6 spinal levels to deliver drugs (10 μl) or cells (2 × 10^5^ in 10 μl PBS) using a 30G needle.

### Depletion of sex hormones by ovariectomy (OVX) and orchidectomy (ORX) surgeries

Animals were anaesthetized with isoflurane for both surgeries (Luo et al., 2021). For conducting OVX, a single midline dorsal incision of 0.5 cm was made to penetrate the skin. Subcutaneous connective tissue was gently freed from the underlying muscle on both sides using blunt forceps. A small incision (<1 cm) was made on each side to gain entry to the peritoneal cavity. Ovary under the thin muscle layer was located and exposed using blunt forceps. Ovary was removed after the oviduct being ligated and severed, and the muscle layer and skin incision were closed by suture. For performing ORX, a single incision was made on the ventral side of the scrotum. The cremaster muscles were cut, and the testicular fat pad was exposed. A single ligature was made around the blood vessels to prevent bleeding following removal of testis from both sides.

### Preparation of BMSC primary cultures

Primary cultures of BMSCs were prepared from CD1 donor mice (same age as receiver mice) under aseptic conditions (Chen et al., 2015). After the mice were sacrificed, both ends of the tibiae and femurs were cut off by scissors, and a syringe fitted with 20-gauge needle was inserted into the shaft of the bone to flush out bone marrow in a modified Eagle’s medium. The cells were cultured in 75 cm^2^ flasks or 6-well culture plates. The property of expanded cells was validated in by flow cytometry using monoclonal antibodies against CD45 (1:400, eBioscience) and CD90 (1:400, eBioscience), as reported (Chen et al., 2015). Cells typically reached confluence in 10-12 days (Figure 1).

**Figure 1.**
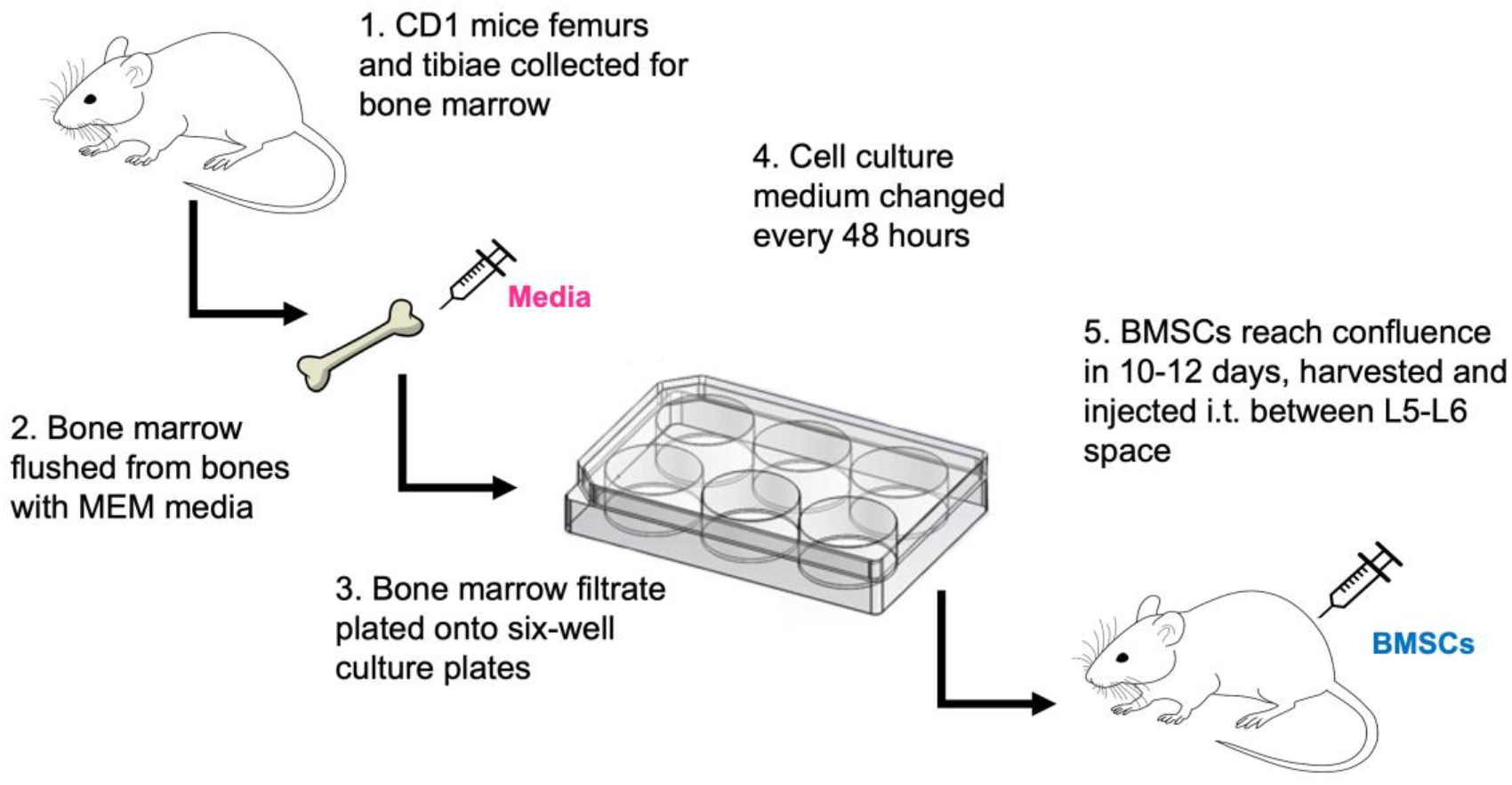
Experimental design showing tissue collection and tissue culture of BMSCs and adoptive transfer of BMSCs to mice via intrathecal injection.

### Behavioral analysis

For von Frey testing for mechanical pain, animals were habituated in boxes on an elevated metal mesh floor under stable room temperature and humidity at least 2 days prior to experiments. Logarithmically graded von Frey fibers (0.02-2.56 gram, Stoelting) were applied to the plantar surface of hindpaw. Paw withdrawal threshold was calculated using the up-down method (Luo et al., 2018). Paw withdrawal frequency was assessed by applying a 0.4 gram von Frey filament to the plantar surface of the hind-paw ten times. Hot plate testing was performed to assess heat sensitivity with a measurement cutoff of 40 seconds (Bioseb in Vivo Research Instrument) following the previous protocol (Han et al., 2016). Cold sensitivity was evaluated by acetone test. We gently applied 20 μl of acetone to the hindpaw bottom using a pipette and scored the responses as: 0, no response; 1, quick withdrawal, paw stamping or flicking; 2, prolonged withdrawal or repeated flicking of the paw; 3, repeated paw flicking and licking. The experimenters were blinded to the treatments. Animals were randomly assigned to each experimental group. The sample sizes were justified based on our previous studies comparing sex difference in pain (Luo et al., 2019, 2021).

### ELISA

TGF-β1 ELISA was performed using culture medium of BMSCs prepared from male and female mice. For each assay, 50 μl of culture media were used and ELISA was conducted according to manufacturer’s instructions. The standard curve was included in each experiment.

### Patch clamp recordings in spinal cord slices

A portion of the lumbar spinal cord (L4-L5) was removed from male and female mice of 6-8 weeks old under urethane anesthesia (1.5 - 2.0 g/kg, i.p.) and kept in pre-oxygenated ice-cold Krebs solution (Wang et al., 2020). Transverse spinal cord slices (400 μm) were cut on a vibrating microslicer. The slices were perfused with Kreb’s solution for 1-3 h prior to experiment. The Kreb’s solution contains (in mM: NaCl 117, KCl 3.6, CaCl2 2.5, MgCl2 1.2, NaH2PO4 1.2, NaHCO3 25, and glucose 11) and was saturated with 95% O2 and 5% CO2 at 36°C. The whole-cell patch-clamp recordings were made in voltage-clamp mode from spinal cord lamina IIo neurons. Majority of these interneurons express somatostatin and are excitatory. They receive synaptic connections from C-fiber afferents and send outputs to lamina I projection neurons (Todd, 2010; Braz et al., 2014; Liu et al., 2016; Chamessian et al., 2018; Duan et al., 2018). Patch pipettes (4-6 MΩ resistance) were pulled from borosilicate capillaries using a P-97 Flaming/Brown micropipette puller (Sutter Instrument Co.). For recording sEPSCs, lamina II neurons were held at holding potentials at −70 mV after establishing the whole-cell configuration. A typical patch pipette had resistance of 5-10 MΩ. We used the following internal solution (in mM: 135 potassium gluconate, 5 KCl, 0.5 CaCl2, 2 MgCl2, 5 EGTA, 5 HEPES, and 5 ATP-Mg). Membrane currents were amplified in voltage-clamp using an Axopatch 200B amplifier (Molecular Devices) and a Digidata 1440A (Axon Instruments). Signals were filtered at 2 kHz and digitized at 10 kHz. Data were stored with a personal computer using pClamp 10 software and analyzed with Mini Analysis (Synaptosoft).

### Statistics

All data were expressed as the mean ± SEM. The sample size for each experiment is indicated in the figure legends. The data were analyzed student t-test (two group comparison) and Two-Way or One-Way ANOVA, followed by Bonferroni’s post-hoc test. A p-value of p<0.05 was considered as statistically significant.

## Results

### Intrathecal injection of BMSCs produces sex-dependent reduction of mechanical pain after CIPN

Our previous study showed that a single intrathecal injection of BMSCs (2.5×10^5^ cells), given 4 days after nerve injury, completely prevented the development of nerve injury-induced neuropathic pain in male mice (Chen et al., 2015). As the first step, we asked whether this approach could also prevent CIPN-induced neuropathic pain. Following four injections of PTX on Days 1, 3, 5, and 7, we gave a single injection of BMSCs (2.5×10^5^ cells) on day 10 (day 4 after the PTX treatment) (Fig. 2).

**Figure 2.**
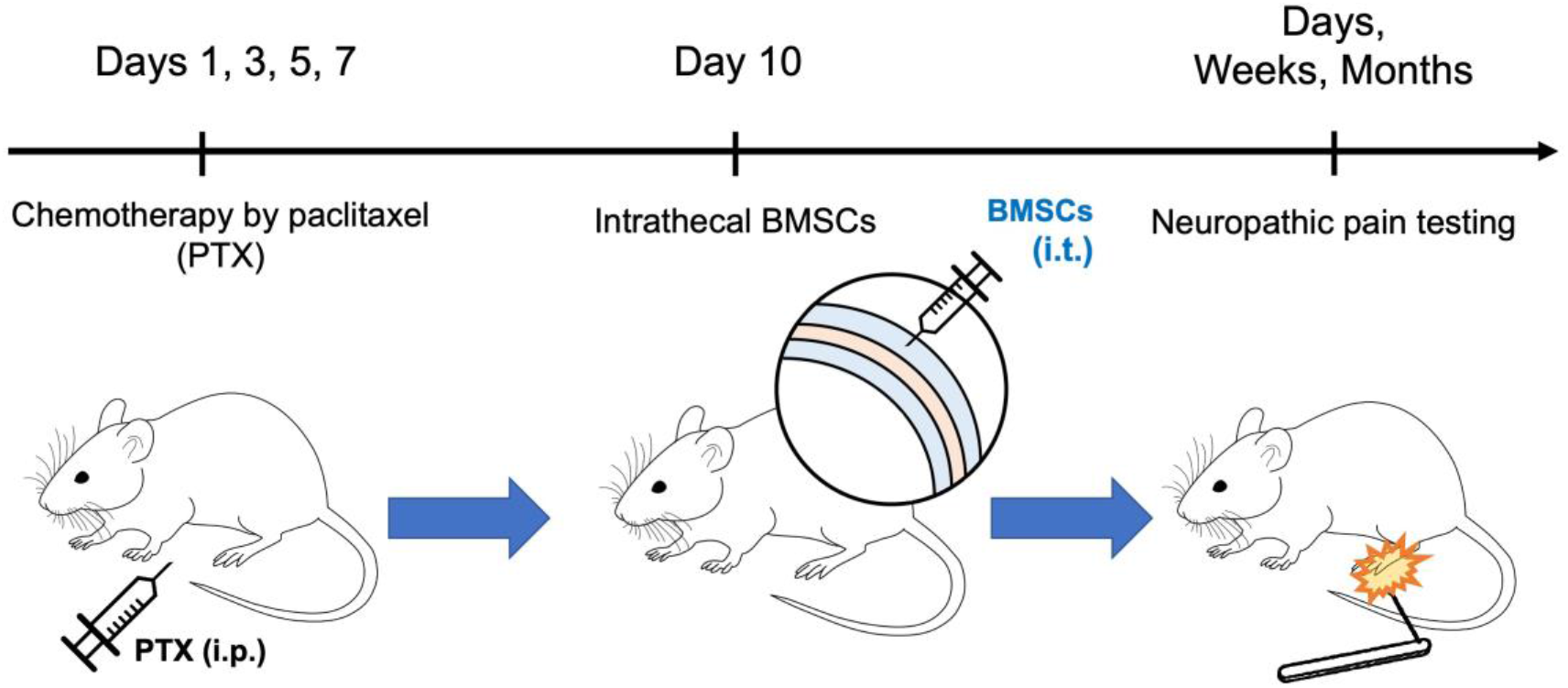
Experimental paradigm showing the generation of CIPN pain model, intrathecal treatment of BMSCs, and behavioral testing for mechanical pain. Notably, 4 injections of PTX were given on Days 1, 3, 5, and 7 and a single injection of BMSCs (2.5×10^5^ cells) was given on day 10. Mechanical pain was assessed by von Frey filament.

In vehicle treated animals, PTX evoked robust mechanical allodynia, which was evident from week 1 to week 7 after the PTX treatment. There was a tendency toward to recovery on week 8 (Fig. 3). Strikingly, a single injection of BMSCs was sufficient to produce long-lasting and significant anti-allodynic effect (p<0.05) for more than 7 weeks in male mice (Fig. 3). We did not see significant effects of BMSCs on cold allodynia in the acetone test (data not shown). Thus, we focused on mechanical pain in this study.

**Figure 3.**
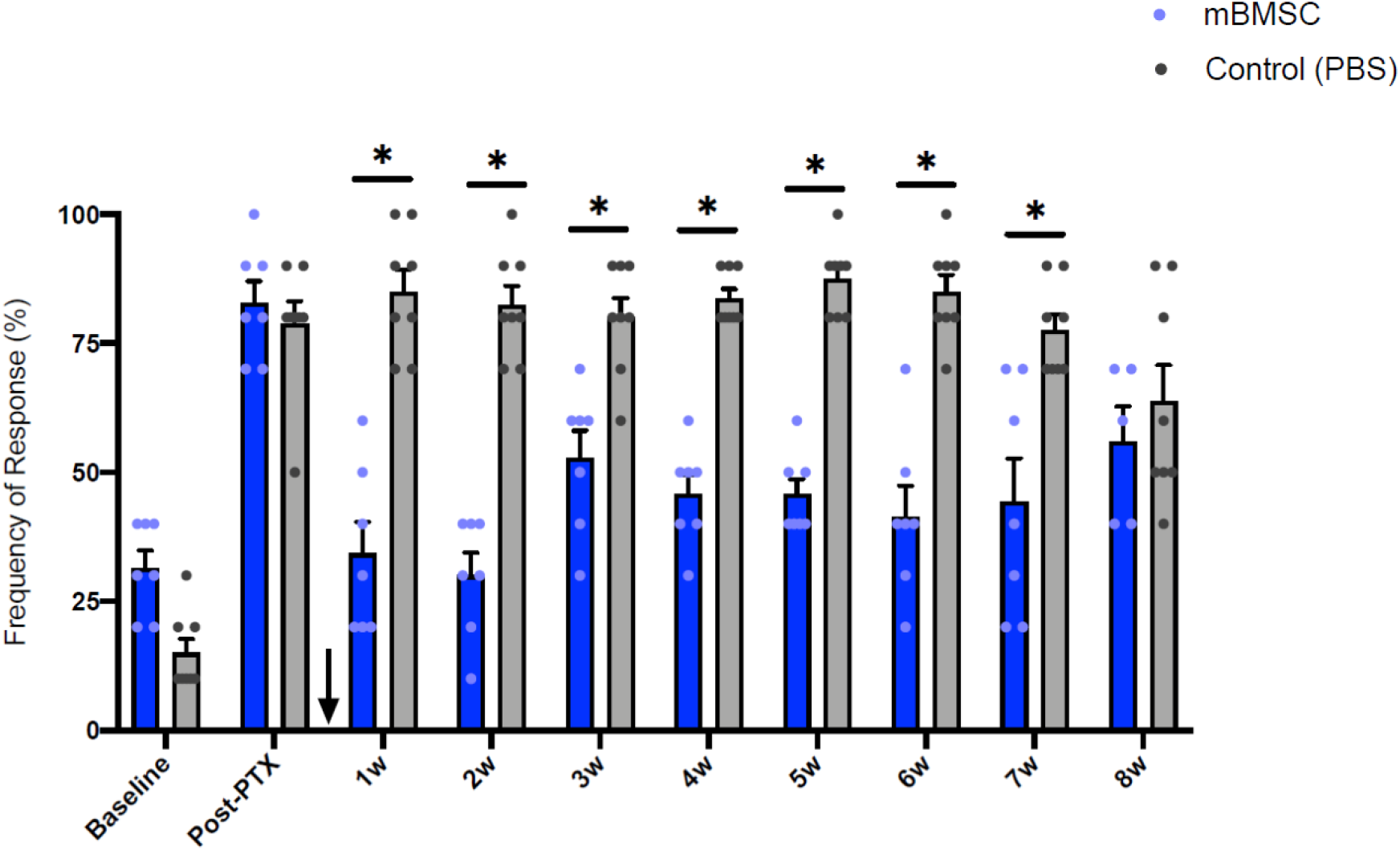
Intrathecal injection of mouse BMSCs (mBMSCs) is sufficient prevent paclitaxel (PTX)-induced mechanical allodynia in male mice. A single injection of BMSCs was given at 4 days post the last PTX injection, i.e.10 days post the first PTX injection (indicated by arrow). Note the long-lasting anti-allodynic effect of BMSCs for more than 7 weeks after a single injection. \**p*<0.05 in comparison to vehicle (PBS). Two-way ANOVA with Bonferroni’s post-hoc test, n=5 mice per group. Data are expressed as the mean ± SEM.

Next, we investigated whether same sex or cross sex transfer of BMSCs might produce different pain control in male and female mice, as shown in the experimental paradigm (Fig. 4A). We tested four different treatments: 1) male BMSCs to male receivers (Fig. 4B), 2) female BMSCs to male receivers (Fig. 4B), 3) male BMSCs to female receivers (Fig. 4C), and 4) female BMSCs to female receivers (Fig 4C). Surprisingly, only male-male transfer of BMSCs exhibited significant analgesic action (p<0.05, Fig. 4B). But males did not show response to BMSCs from female donors (Fig. 4B). Female mice responded to neither male BMSC donors nor female BMSC donors (Fig. 4C). Together, these findings suggest a strong sex dimorphism in BMSC-induced pain relief.

**Figure 4.**
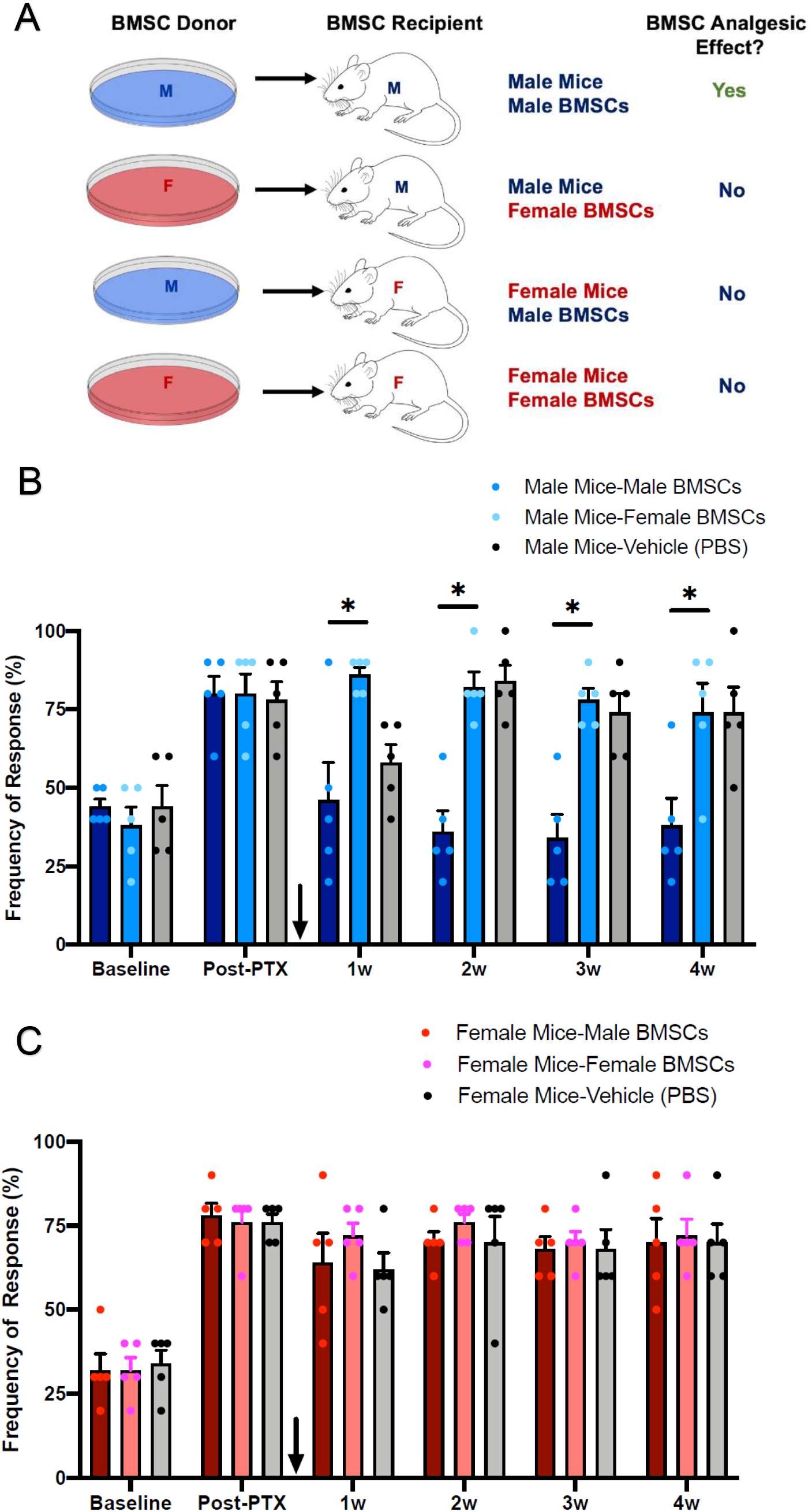
Effects of same sex and cross sex transfer of BMSCs on chemotherapy-induced mechanical allodynia in male and female mice. (A) Experimental paradigm showing same sex and cross sex transfer of BMSCs and summary of analgesic actions. Note only male-male transfer shows analgesic action. (B, C) Effects of same sex and cross sex transfer of BMSCs on chemotherapy-induced mechanical allodynia in male (B) and female (C) receivers. \**p*<0.05, male vs. female BMSCs. Two-way ANOVA with Bonferroni’s post-hoc test, n=5 mice per group. Arrow indicates the time of BMSCs injection. Data are expressed as the mean ± SEM.

### TGF-β1 mediates BMSC-induced analgesia and synaptic modulation in males

Intrathecal injection of low-dose of TGF-β1 has been shown to reduce neuropathic pain in male rats and mice with nerve injury (Echeverry et al., 2009; Chen et al., 2015). To determine the mechanism by which BMSCs would inhibit pain, we tested the involvement of TGF-β1 signaling, which was required for BMSC-induced pain in the nerve trauma model (Chen et al., 2015). Notably, BMSC-induced analgesia was reversed by intrathecal injection of anti-TGF-β1 antibody (10 μg) in male mice at 3h after the injection: no statistical difference was observed between the vehicle-treated and BMSC-treated animals at this time point (Fig. 5A). ELISA analysis detected TGF-β1 secretion in culture medium of BMSCs from male and female donors. However, TGF-β1 secretion was significantly higher in BMSCs from male than female donors (*p*<0.05, Fig. 5B).

**Figure 5.**
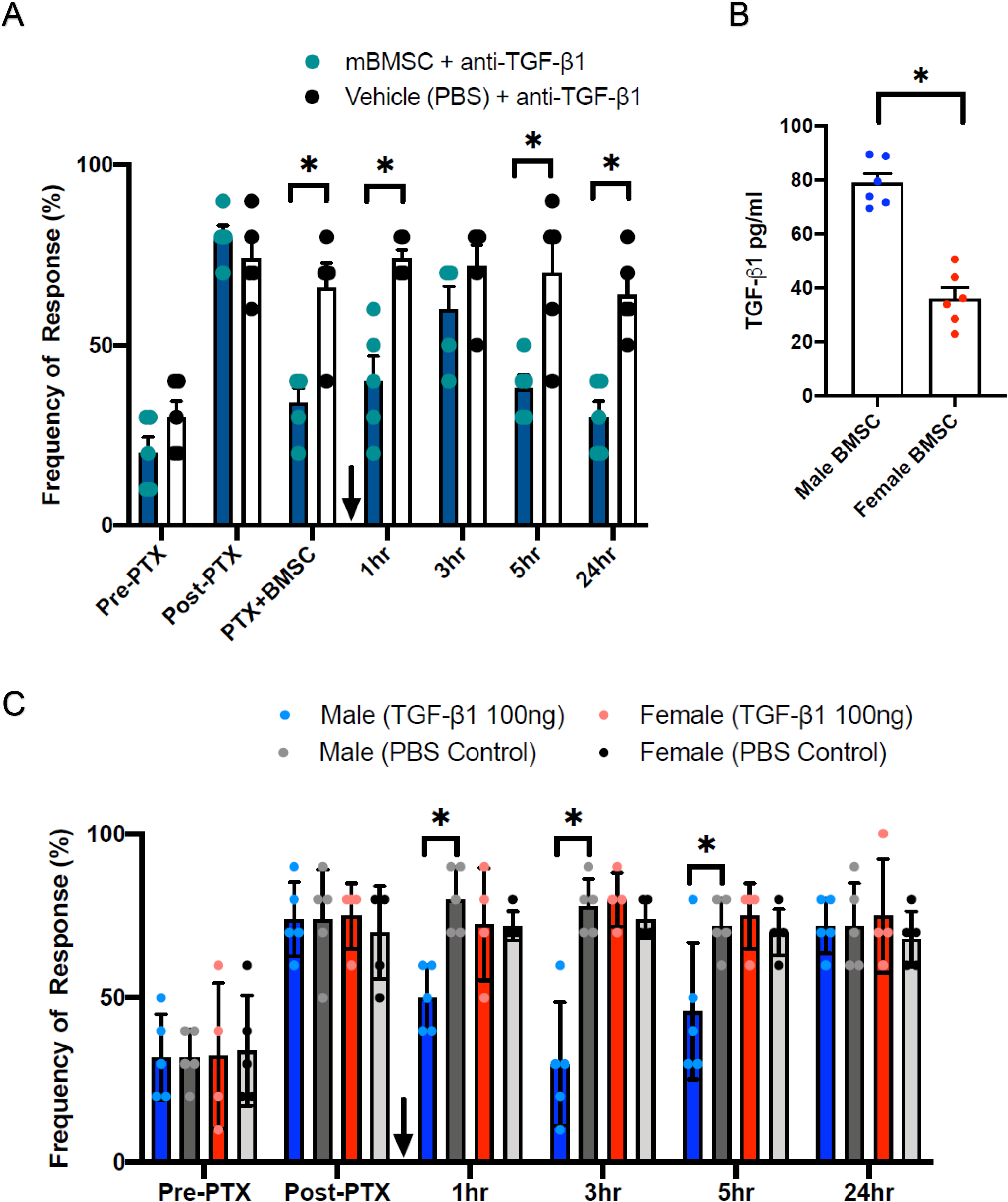
TGF-β1 mediates BMSC-induced analgesia and sex dimorphism following chemotherapy. (A) Reversal of BMSC-induced analgesia by intrathecal injection of anti-TGF-β1 antibody (10 μg) in male mice. \**p*<0.05, BMSC vs. vehicle treated animals. Two-way ANOVA with Bonferroni’s post-hoc test, n=5 mice per group. Note the analgesic action (anti-allodynia) is lost 3 hours after the antibody treatment. (B) ELISA analysis showing TGF-β1 secretion in culture medium of BMSCs from male and female donors. \**p*<0.05, unpaired t-test, n=6 separate cultures. (C) Intrathecal injection of TGF-β1 (100 ng) reduced PTX-induced mechanical allodynia in male but not female mice. \**p*<0.05 in comparison to vehicle (PBS) in males. Two-way ANOVA with Bonferroni’s post-hoc test, n=5 mice per group. Note a single injection of TGF-β1 produces significant inhibition of mechanical allodynia at 1h, 3h, and 5h. Arrow indicates the antibody injection. Data are expressed as the mean ± SEM.

Next, we examined whether exogenous TGF-β1 would affect neuropathic pain in either sex. Strikingly, intrathecal injection of TGF-β1 (100 ng) reduced PTX-induced mechanical allodynia only in male mice, but not female mice. A single injection of TGF-β1 elicited significant inhibition of mechanical allodynia at 1h, 3h, and 5h in males, but mechanical pain returned at 24h (Fig. 5C).

### Exogenous TGF-β1 rapidly suppresses chemotherapy-induced synaptic plasticity in spinal cord neurons of male not female mice

Synaptic plasticity plays a crucial role in inflammatory pain and neuropathic pain (Woolf and Salter, 2000; Gruber-Schoffnegger et al., 2013; Li et al., 2014; Stockstill et al., 2018). We recorded spontaneous excitatory postsynaptic currents (sEPSCs) in lamina IIo neurons of spinal cord slices from control male mice, female mice injected with PTX, and male mice injured with PTX. We then perfused spinal cord slices with TGF-β1 (60 ng/ml) and tested its actions on sEPSCs. *In vivo* PTX treatment increased sEPSC frequency in lamina IIo neurons recorded ex vivo (Fig. 6A,B). Perfusion of TGF-β1 had no effects on sEPSCs in neurons of female mice but reduced sEPSC in neurons of male mice. A significant difference was found between male and female neurons with PTX treatment (p<0.05, Fig. 6A,B). Notably, TGF-β1 had no effects on sEPSCs amplitude (Fig. 6C). Thus, it is plausible that TGF-β1 may inhibit sEPSCs through presynaptic modulation.

**Figure 6.**
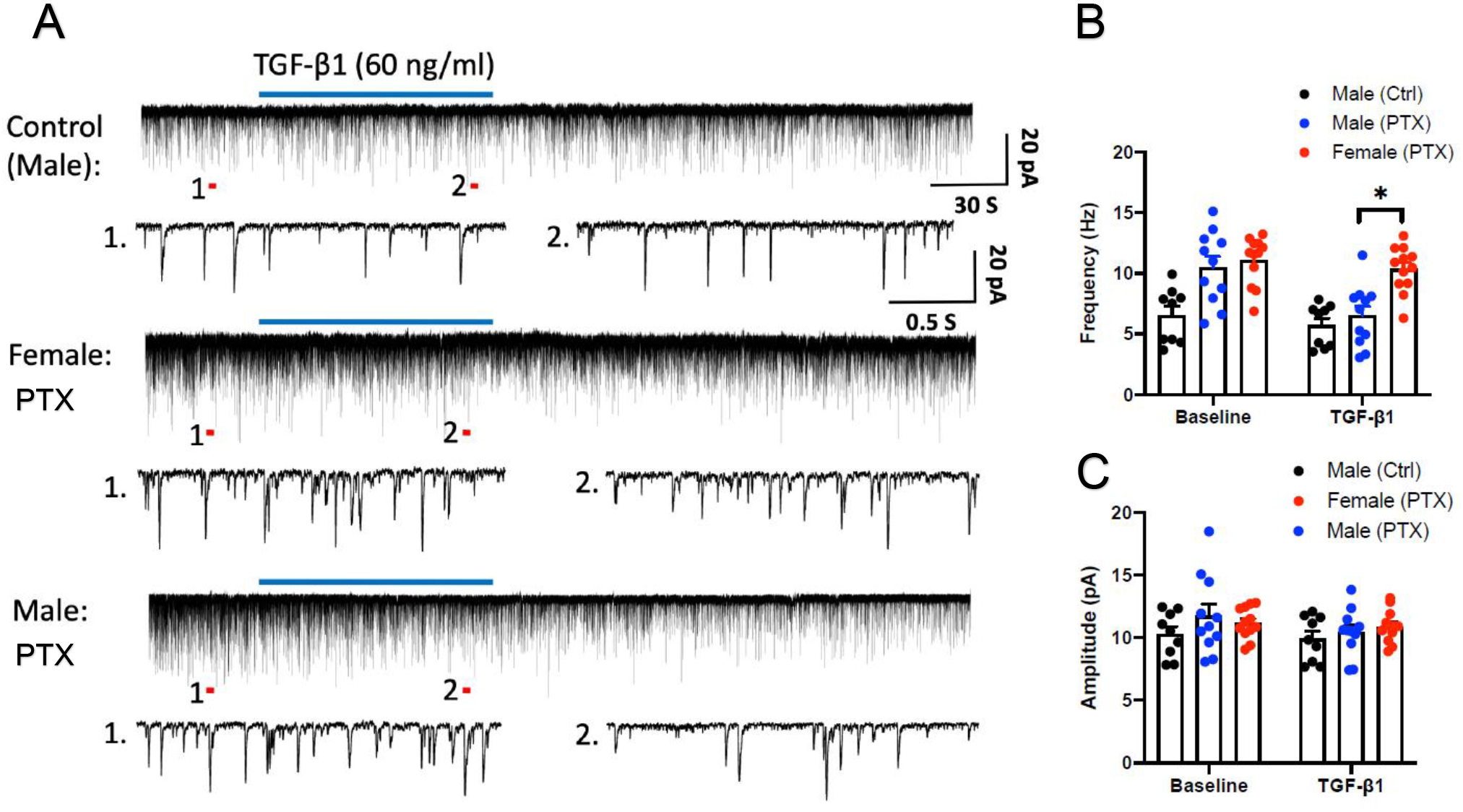
Exogenous TGF-β1 rapidly suppresses PTX-induced enhancement of excitatory synaptic transmission in lamina IIo neurons of spinal cord slices of male not female mice. (A) Traces of spontaneous excitatory postsynaptic currents (sEPSCs) in lamina IIo neurons of spinal cord slices from control male mice (top row), female mice with PTX (middle row), and male mice with PTX (bottom row). Note sEPSCs are increased after PTX chemotherapy treatment. Also note that TGF-β1 (60 ng/ml) perfusion inhibited sEPSCs in male but not female slices. Trace 1 and 2 are enlarged in each row. (B, C) Quantification of sEPSC frequency (B) and amplitude (C). \**p*<0.05, male vs. female in the presence of PTX and TGF-β1. Note that TGF-β1 has no effects on the amplitudes of sEPSCs in both sexes. n=9-12 neurons/group. Statistical significance was determined by one-way ANOVA followed by Bonferroni post-hoc test. All data are expressed as mean ± S.E.M.

### Exogenous TGF-β1 has distinct effects on mechanical, cold, and heat sensitivity in naïve animals

To further investigate the role of TGF-β1 in homeostatic regulation of pain, we tested the effects of TGF-β1 on different modalities of pain in naïve animals. Paw withdrawal threshold analysis showed that intrathecal TGF-β1 (100 ng) caused a transient increase in withdrawal threshold in males (p<0.05, at 0.5h, Fig. 7A) but failed to do so in females (Fig. 7B). It is suggested that TGF-β1 decreased mechanical sensitivity in males even in hemostatic physiological conditions. Interestingly, acetone testing revealed significant increase in cold response score in males (p<0.05 at 0.5h and 1h, Fig. 7C) and females (p<0.05 at 0.5h, Fig. 7D) following intrathecal TGF-β1 (100 ng). Hot plate testing showed unaltered heat pain sensitivity following intrathecal administration of TGF-β1 (100 ng) in males (Fig. 7E) and females (Fig. 7F). Together, these results suggest 1) TGF-β1 regulates pain under the hemostasis condition; 2) TGF-β1 differentially regulates mechanical pain, cold pain, and heat pain; and 3) TGF-β1-induced sex dimorphism mainly manifests in mechanical pain.

**Figure 7.**
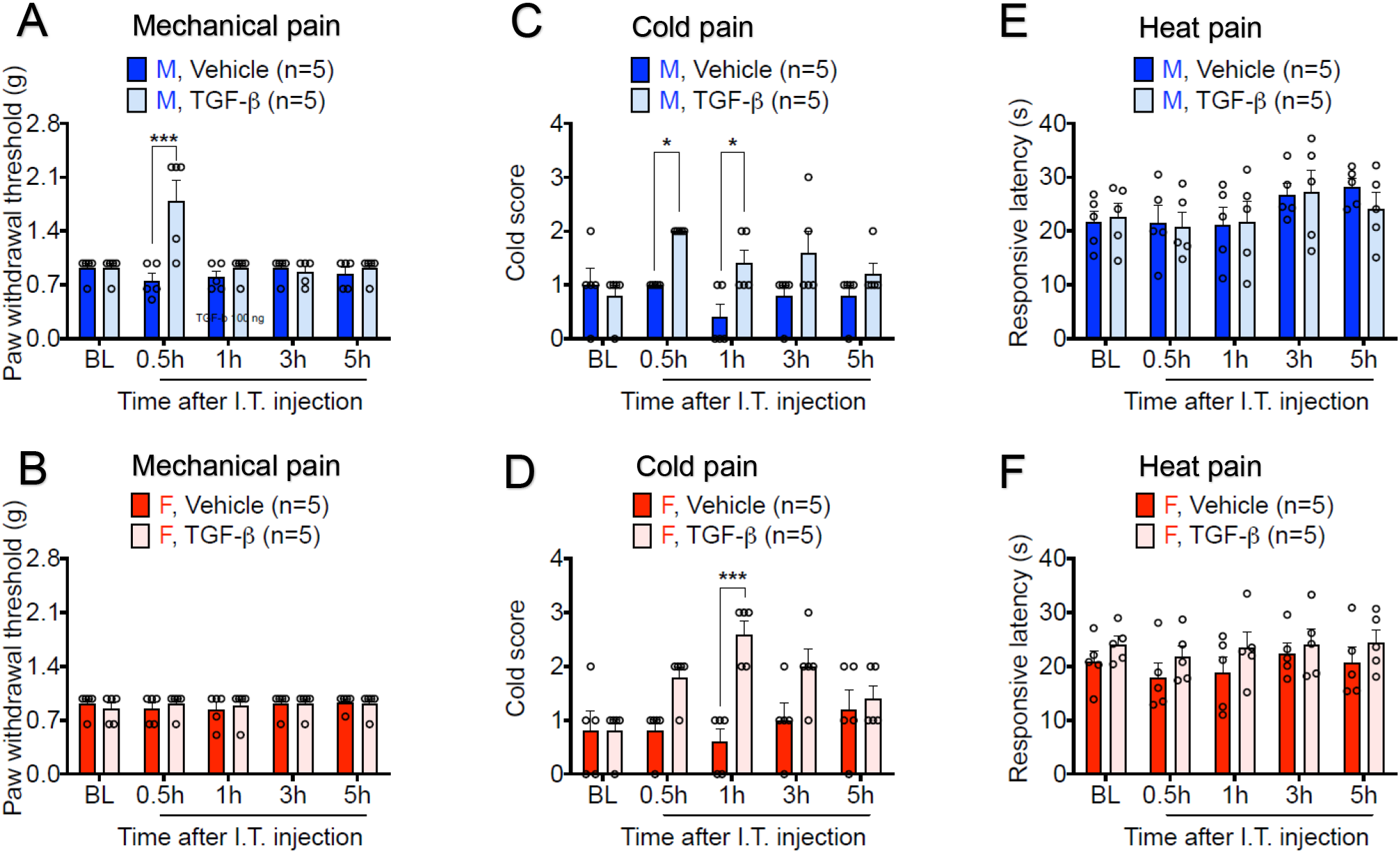
Distinct effects of intrathecal injection of TGF-β1 (100 ng) on mechanical, cold, and heat pain sensitivity in naïve animals of males and females. (A, B) von Frey testing showing paw withdrawal threshold in males (A) and females (B). \*\*\**p*<0.001 in comparison to vehicle (PBS) in males. (C, D) Acetone testing showing cold response score in males (C) and females (D). \**p*<0.05, \*\*\**p*<0.001, in comparison to vehicle in males and females. (E, F) Hot plate testing showing heat pain sensitivity (response latency) in males (E) and females (F). Two-way ANOVA with Bonferroni’s post-hoc test, n=5 mice per sex per group. Note that TGF-β1 decreases mechanical sensitivity in males, increases cold sensitivity in both sexes, and has no effects on heat pain sensitivity in neither sex. Data are expressed as the mean ± SEM.

### TGF-β1 modulation of mechanical pain depends on sex hormones

It was widely believed that sex hormones mediate sex dimorphism in pain (Hucho and Levine, 2007; Luo et al., 2021). To this end, we conducted orchidectomy (ORX) and ovariectomy (OVX) surgeries to deplete sex hormones in males and females, respectively. We then intrathecally injected TGF-β1 (100 ng) in males with ORX and in females with OVX, using sham surgery as control. Of interest TGF-β1 increased paw withdrawal threshold in males with sham surgery but not in males with ORX (p<0.05 at 0.5h, Fig. 8A). Therefore, male hormones are required for TGF-β1 to inhibit mechanical pain in male mice. In contrast, TGF-β1 failed to change paw withdrawal threshold in females with sham surgery or females with OVX (Fig. 8B), suggesting that female hormones may not be required to regulate TGF-β1-medidated pain.

**Figure 8.**
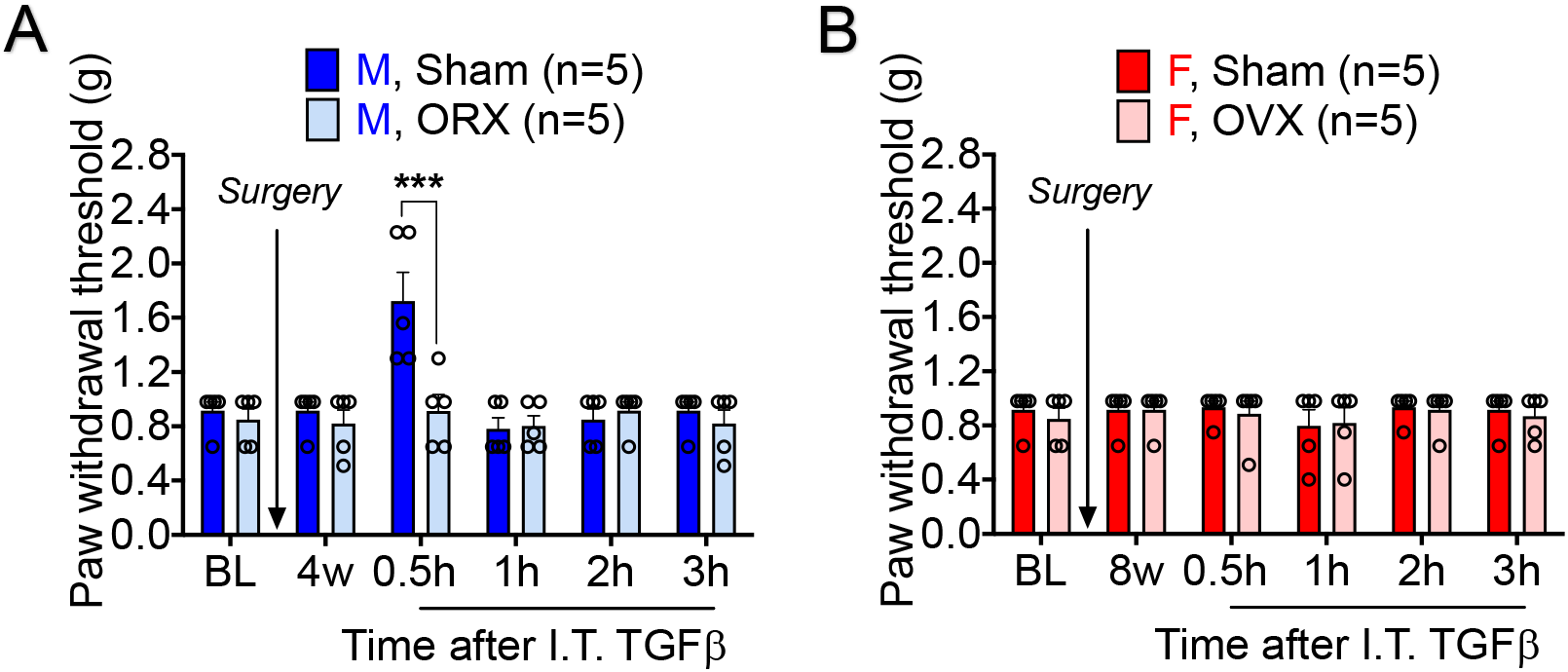
Effects of orchiectomy (ORX) and ovariectomy (OVX) surgery and TGF-β1 on mechanical pain by von Frey testing. (A) Paw withdrawal threshold in male mice with ORX and sham surgery. (B) Paw withdrawal threshold in female mice with OVX and sham surgery. All mice were given intrathecal injection of 100 ng of TGF-β1. ***p<0.001, Two-way ANOVA with Bonferroni’s post-hoc test, n=5 mice per sex per group. Note that TGF-β1 decreases mechanical sensitivity only in males with sham surgery. Data are expressed as the mean ± SEM.

## Discussion

Recent progress has demonstrated sex dimorphism in pain regulation by several immune and glial cell types, including microglia (Sorge et al., 2015; Taves et al., 2016; Mapplebeck et al., 2018), macrophages (Luo et al., 2019; Yu et al., 2020; Luo et al., 2021), and T cells (Sorge et al., 2015). Systemic and local administration of MSCs/BMSCs has demonstrated long-term analgesic potency in inflammatory pain, neuropathic pain, and opioid-induced anti-nociceptive tolerance/hyperalgesia (Guo et al., 2011; Chen et al., 2015; Hua et al., 2016; Huh et al., 2017; Buchheit et al., 2020). However, sex dimorphism of pain relief by BMSCs was not investigated in these studies. In the present study, our findings showed that a single intrathecal injection of BMSCs, prepared from male donors, completely prevented the development of chemotherapy-evoked mechanical allodynia in male mice. However, female mice showed no response to either male or female BMSCs. Furthermore, male mice failed to demonstrate an analgesic response to BMSCs from female donors.

Mechanistically, we have provided several lines of evidence to support the involvement of TGF-β1 signaling in BMSC-induced pain inhibition in males. First, BMSCs from males secreted greater amount of TGF-β1 in the culture media, compared to BMSCs from females. Second, BMSC-induced analgesia in males was reversed by intrathecal injection of the anti-TGF-β1 antibody, suggesting that endogenous TGF-β1 signaling is required for the benefit of BMSCs. Third, intrathecal injection of TGF-β1 reverse chemotherapy-induced mechanical allodynia in male mice, in further support of the reports that exogenous TGF-β1 is sufficient to alleviate neuropathic pain in male mice (Echeverry et al., 2009; Chen et al., 2015). Suppression of neuropathic pain by TGF-β1 in male rodents was further demonstrated by the finding that miR-30c-5p drives neuropathic pain via suppressing TGF-β1 expression in male rats (Tramullas et al., 2018). However, the role of TGF-β1 was not examined in female mice in previous studies (Echeverry et al., 2009; Chen et al., 2015; Tramullas et al., 2018). Notably, intrathecal TGF-β1 (100 ng) had no significant effects on chemotherapy-induced mechanical allodynia in female mice.

We also revealed TGF-β1-mediated sex dimorphism can further manifest ex vivo in isolated spinal cord slice preparation. Patch-clamp recordings in lamina II neurons showed that perfusion of TGF-β1 only inhibited neuropathic pain-associated synaptic plasticity (sEPSC frequency increase by paclitaxel) in spinal cord neurons from male mice but showing no effect in neurons of female mice. This result suggested that sex dimorphism may retain at the spinal cord microenvironment, which contains neurons, glial cells, and immune cells. Our previous study showed that the analgesic effect of TGF-β1 is mediated by TGF-β receptor 1 (TGFβR1, encoded by *Tgfbr1*), as SB431542, a potent and selective inhibitor of TGFβR1, could eliminate the analgesic effects of TGF-β1 (Chen et al., 2015). Single-cell RNA sequence analysis revealed *Tgfbr1* expression in primary sensory neurons of DRG, including nociceptors (Usoskin et al., 2015; Renthal et al., 2020). Furthermore, non-neuronal cells, including immune and glial cells express *Tgfbr1*, *Tgfbr2, Tgfbr3*, as well as *Tgfb1* (Renthal et al., 2020). Future studies are needed to investigate sex differences in the expression of these transcripts in DRG and spinal cord under the normal and pathological conditions.

Sex hormones have been strongly implied in sex dimorphism in pain (Dina et al., 2001; Hucho and Levine, 2007; Sorge et al., 2015; Luo et al., 2021). We conducted orchidectomy and ovariectomy surgeries to determine the contributions of male and female hormones to TGF-β1-evoked suppression of mechanical pain. We found that intrathecal TGF-β1 increased paw withdrawal in male mice with sham surgery but failed to do in mice with orchiectomy. This result indicated that male hormones are required for TGF-β1 to inhibit mechanical pain in male mice. By contrast, TGF-β1 did not alter paw withdrawal threshold in female mice with ovariectomy or sham surgery. Thus, female hormones are not critically involved in TGF-β1-medidated pain relief, although estrogen has been shown to promote pain in females (Patil et al., 2019; Luo et al., 2021).

It is noteworthy that some studies have shown analgesic (anti-allodynic) effects of MSCs in both sexes. For example, nasal MSC application was shown to reverse chemotherapy-induced allodynia in male and female mice. Moreover, nasally administered MSC enter the meninges of the brain, spinal cord and peripheral lymph nodes to promote IL-10 production through macrophages (Boukelmoune et al., 2021). This discrepancy may be due to different source of BMSCs/MSCs (cell line vs. primary cultures), different injection routes (nasal vs. intrathecal), different destination of the injected cells (brain vs. DRG), and distinct signaling mechanisms (IL-10 vs. TGF-β1). It is also plausible that sex dimorphism may be partially related to environments in which animals are raised in different animal facilities with distinct food and microbiota composition.

In summary, we have demonstrated a previously unrecognized mechanism of sex dimorphism in pain modulation, by which intrathecally injected BMSCs and TGF-β1 reduced mechanical pain in male mice. Our study has provided new cellular and molecular insights into sex dimorphism in pathological conditions (chemotherapy) and physiological conditions (naïve animals). Recently, we proposed that crosstalk between immune cells and sensory neurons may underlie sex dimorphism in pathological pain conditions, such as CIPN (Luo et al., 2021). It will be of great interest to investigate sex dimorphism in TGF-β1/TGFBR signaling in immune cells (including BMSCs) and glial cells and their interactions with primary sensory neurons during the pathogenesis and resolution of pain.

## Acknowledgements

This work was supported by Duke University Anesthesiology Research Funds and partially supported by NIH grants R01-DE17794 and DoD grants W81XWH2110885 and W81XWH2110756.

## Notes

**Conflict of interest statement**: All the authors have no competing financial interest in this study.

### Competing Interest Statement

The authors have declared no competing interest.

